# Visualization of axonal protein allocation in *Drosophila* with whole-brain localization microscopy

**DOI:** 10.1101/454942

**Authors:** Li-An Chu, Chieh-Han Lu, Shun-Min Yang, Kuan-Lin Feng, Yen-Ting Liu, Chun-Chao Chen, Yun-Chi Tsai, Peilin Chen, Ting-Kuo Lee, Yeu-Kuang Hwu, BiChang Chen, Ann-Shyn Chiang

**Author notes:** These authors contributed equally to the project (Li-An Chu, Chieh-Han Lu). Present address: Department of Genetics and Complex Diseases, Harvard T H Chan School of Public Health, Boston, MA02115, United States (Chieh-Han Lu). Correspondence and requests for materials should be addressed to A.S.C. or B.C.C Ann-Shyn Chiang, Brain Research Center, National Tsing Hua University, 101, Section 2, Kuang-Fu Road, Hsinchu 30013, Taiwan; Bi-Chang Chen, Research Center for Applied Sciences, Academia Sinica, 128 Sec. 2, Academia Rd., Nankang, Taipei 11529, Taiwan.

## Abstract

Long-term memory (LTM) formation requires learning-induced protein synthesis in specific neurons and synapses within a neural circuit. Precisely how neural activity allocates new proteins to specific synaptic ensembles, however, remains unknown. We developed a deep-tissue super-resolution imaging tool suitable for single-molecule localization in intact adult *Drosophila* brain, and focused on the axonal protein allocation in mushroom body (MB), a central neuronal structure involved in olfactory memory formation. We found that insufficient training suppresses LTM formation by inducing the synthesis of vesicular monoamine transporter (VMAT) proteins within a dorsal paired medial (DPM) neuron, which innervates all axonal lobes of the MB. Surprisingly, using our localization microscopy, we found that these learning-induced proteins are distributed only in a subset of DPM axons in specific sectors along the MB lobes. This neural architecture suggests that sector-specific modulation of neural activity from MB neurons gates consolidation of early transient memory into LTM.

## Introduction

Memory formation requires learning-induced protein synthesis in specific neurons and synapses within a neural circuit. In all species examined, there are two distinct phases of memory formation: a transient neural activity associated with early memory and a protein-synthesis-dependent change in synaptic connectivity associated with long-term memory (LTM)^1,2^. Early memory is labile; sustained neural activity during this phase nonetheless is crucial for the induction of LTM and its underlying protein synthesis, which occurs only in a few neurons sparsely distributed throughout the brain^3–5^. Exactly how neural activity induces protein synthesis in some but not all neurons in a circuit and then allocates new proteins to specific synaptic ensembles during LTM formation, however, remains unknown.

Recent advances in super-resolution microscopy have allowed for localization of single molecules within individual cells^6–8^, but not within large tissues^9^. Our understanding of memory formation may benefit from these advances by enabling us to visualize the allocation of associative learning-induced proteins at the level of the synapse. In the present study, we integrated several optical technologies to develop a deep-tissue imaging tool suitable for single-molecule localization in an intact adult *Drosophila* brain. We show that insufficient training suppresses LTM formation by inducing the synthesis of vesicular monoamine transporter (VMAT) proteins in a single dorsal paired medial (DPM) neuron, which innervates all axonal lobes of the mushroom body (MB). Consistent with this observation, downregulation of VMAT or reduced serotonin synthesis in the DPM neuron enhances LTM formation. Strikingly, we found that training-induced VMAT proteins are preferentially allocated to a specific subset of DPM neurites, which arborize within the α2/β′1 MB sectors. This neural architecture suggests that sector-specific serotonin modulation of neural activity from MB neurons gates consolidation of early transient memory into LTM. Moreover, the present study demonstrates that our single-molecule imaging technique can be used to visualize memory allocation at specific synaptic ensembles within an intact brain.

## Results

### Deep-tissue localization microscopy (DTLM)

The study of memory formation requires novel tools for visualizing the allocation of learning-induced proteins into synaptic ensembles in an intact brain. Several research groups have attempted to modify super-resolution microscopy techniques for use with larger imaging volumes^10–13^. Among these techniques, point accumulation for imaging in nanoscale topography has achieved sub-100 nm resolution in samples over 20-µm thick by utilizing the inherent optical sectioning of the lattice lightsheet to prevent premature photo-bleaching (which limits localization and image quality)^13,14^. This method nonetheless fails to compensate for tissue-induced aberrations, thereby restricting imaging depth and volume. To visualize protein molecules within larger tissues, we integrated a Bessel beam lightsheet, spontaneous blinking fluorophore, and optical tissue clearing to develop a deep-tissue imaging tool suitable for single-molecule localization in an intact adult *Drosophila* brain (**Fig. 1a and Methods)**. The lightsheet was generated by scanning a Bessel beam created by filtering the laser illumination with an annular ring mask at the Fourier plane, which was conjugated to the back aperture of the excitation objective (customized, N.A. = 0.5, working distance = 12.8 mm; NARLabs, ITRC, Taiwan; **Fig. 1b**)^15,16^. The Bessel beam has been described as a self-reconstruction light beam that is particularly effective for penetrating into a thick specimen^15,16^. The length of this lightsheet was extended from 50 μm to over 200 μm using an axicon lens (**Supplementary Fig. 1a,b**). Through constructive interference at the imaging plane, the energy distribution of the lightsheet is spatially confined within a 0.5-μm layer, thereby reducing background signal and preventing unwanted photo-bleaching (**Supplementary Fig. 1c,d**).

**Fig. 1.**
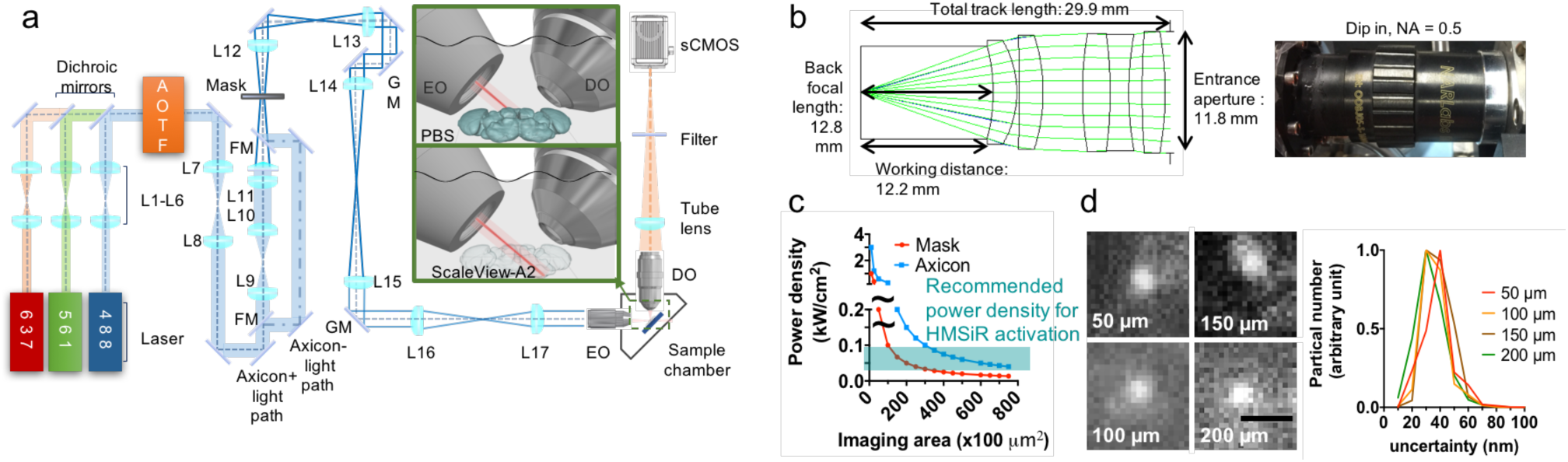
Bessel beam lightsheet for deep-tissue localization microscopy (DTLM). **(a)** The schematic configuration of the DTLM system. Inset: the geometry of relative positions between two objective lenses and the specimen. AOTF: acoustic-optical tuneable filter; EO, excitation objective; DO, detection objective; GM: galvo mirror; L: lens. **(b)** A customized dip-in objective lens matching refractive index of ScaleView-A2 at a long working distance of 12.2 mm. Left, Photograph of the lens. Right, Schematic design and simulated ray tracing. **(c)** The lightsheet generated using an axicon lens carries higher power density than that generated using the mask only. At the power density (40 w/cm^2^) for HMSiR excitation, the maximum lightsheet areas are 25,000 μm^2^ and 75,000 μm^2^, generated using a mask only and an axicon lens, respectively. **(d)** Representative HMSiR blinking events (left) and overall lateral uncertainty distribution (right) at four different depths in the fly brain cleared with ScaleView-A2. Scale bar = 1 μm.

Traditional localization microscopy relies on two chemical mechanisms for the partial activation of fluorophores: either an alternating two-wavelength exposure or a single-wavelength high-intensity illumination^7,8,17^. Use of an additional activation laser, however, prolongs image acquisition time and causes additional photo-bleaching due to short-wavelength exposure. For high intensity illumination, the laser fluence deposited to the sample rapidly consumes the photon budget and makes it unrealistic to achieve large scale imaging. We addressed these problems by using a novel spontaneous blinking fluorophore, HMSiR, which can be excited at a relatively low power density of 40 W/cm^2^. Consequently fluorophore blinking was extended to an area up to 75,000 μm^2^ **(Fig. 1b)**. When combined with the use of HMSiR and tissue clearing (ScaleView-A2)^*18*^, our DTLM method can reconstruct super-resolution images of an entire adult brain **(Methods)**. Importantly, the uncertainty of localized blinking events is similar through the entire z depth (**Fig. 1c**). Typically, over 100 M particles are contained in each of the four sub-volumes used to reconstruct DTLM images of the whole brain (**Supplementary Fig. 1e**), and the total acquisition time is less than one day **(Methods)**. To analyze gigantic datasets of raw images and reconstruct the whole fly brain at super-resolution, we created a parallel computing pipeline based on ThunderSTORM^19^ on a Lustre-backed Torque cluster (**Supplementary Movie 1**, **Supplementary Fig. 2**).

The photoelectric sensors used for conventional fluorescence imaging (green fluorescent protein, GFP) have a limited dynamic range. Thus, it can be difficult to capture structures with weaker GFP signals in a single image (**Supplementary Fig. 5b, left**). Because DTLM localizes individual molecules separately and reconstructs the image based on localization events, it is insensitive to intensity differences, thereby enabling the capture of greater detail in a single image. DTLM, for instance, captured a majority of parallel, densely bundled neural fibers connecting the brain and body (**Supplementary Fig. 5b, middle**). When sufficient localization density was achieved, individual brain-ascending/descending fibers could be digitally segmented (**Supplementary Fig. 5b, right**). Furthermore, DTLM enabled three-dimensional visualization of individual synaptic proteins (Down syndrome cell adhesion molecules) within a single spine-like protrusion in a dendrite of the giant fiber neuron (**Supplementary Fig. 5c**).

### Visualizing fine neurites in the whole brain

Targeted genetic manipulations using the promoter-driven *TH-Gal4* line^20^ have revealed that dopaminergic neurons (DANs) are involved in various brain functions in *Drosophila*, including decision making^21^, arousal^22^, and learning and memory^23^. To obtain a more comprehensive understanding of these neural circuits underlying various behaviors, we used single-molecule DTLM imaging to map the morphology and wiring patterns of all *TH-Gal4* neurons in the adult brain (**Fig. 2**). Due to nominal photo-bleaching and optical clearing, DTLM allowed us to image several overlapped sub-volumes under a high N.A. lens and stitch them into a single big dataset of the entire brain at super-resolution. This large-volume super-resolution map enabled simultaneous visualization of putative dopaminergic neurons labelled with strong GFP signal in the central brain (**Supplementary Fig. 6**), as well as fine neurites labelled with weak GFP signal in the optic lobe (**Fig. 2a)**. Serial optical slices and three-dimensional navigation demonstrated extensive yet separable neurites (**Supplementary Movie 3**). These high-quality images also allowed for digital segmentation of most individual neurons, with nominal axial overlap (**Fig. 2b**). Next, we applied DTLM to visualize the allocation of learning-induced new proteins.

**Fig. 2.**
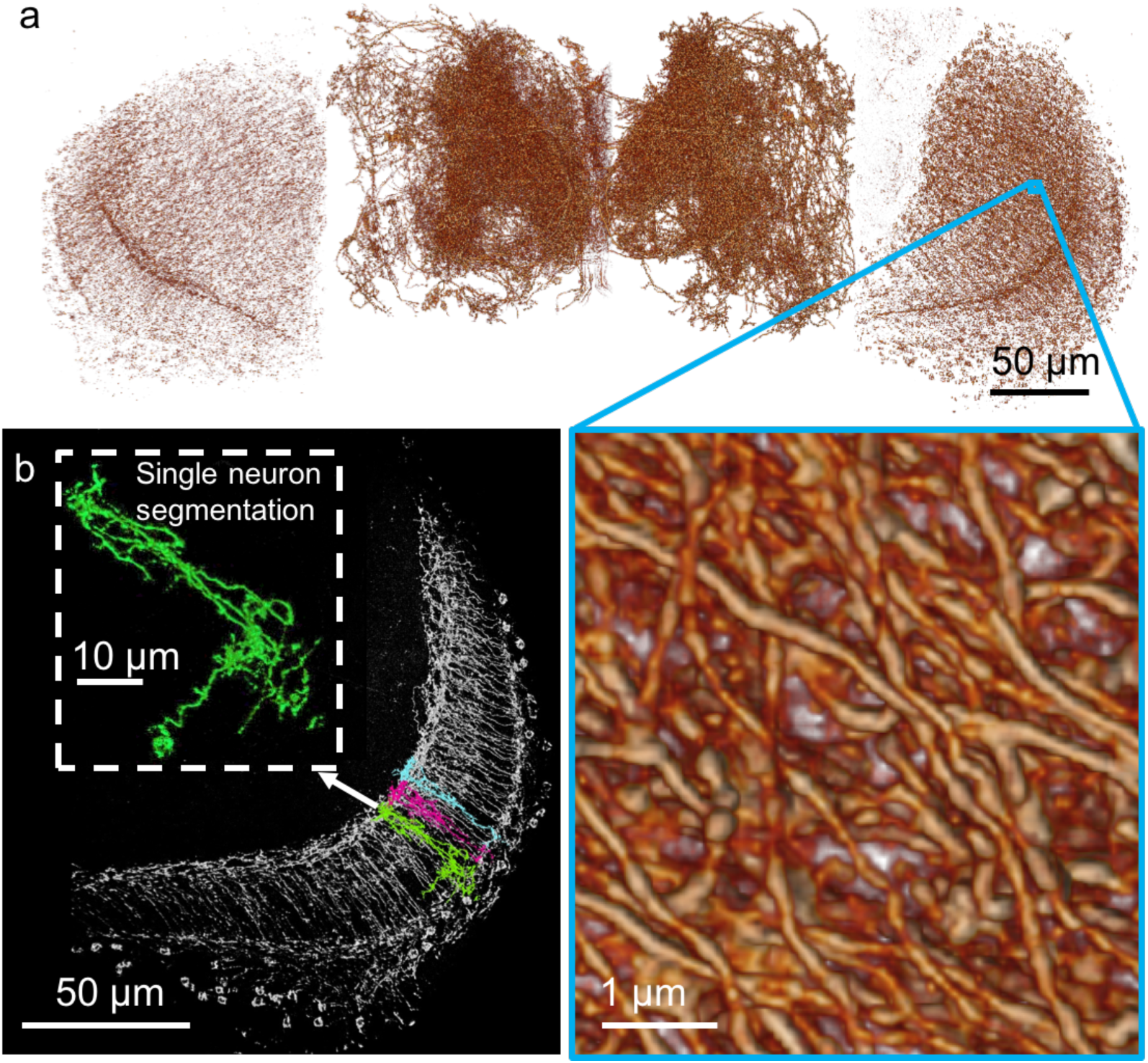
Single-molecule deep-tissue localization microscopy (DTLM) imaging of dopaminergic neurons (DANs) in the whole *Drosophila* brain. **(a)** Volume rendering of whole-brain DANs. Zoom-in images show distinguishable interweaved neurites. **(b)** Digital segmentation of local neurons in the medulla. Inset: enlarged green neuron. The experimental flies carried *TH-Gal4*; *UAS-GCaMP6f* transgenes.

### Suppression of LTM formation due to insufficient training

*Drosophila* can learn to associate an odor with foot-shock punishment, and memory formation thereafter exhibits several temporal phases, including protein synthesis-dependent long-term memory^24,25^. The odor-shock association appears to occur in the MB, where the conditioned odor stimulus is represented by the neural activity of a sparse subpopulation of intrinsic Kenyon cells (KCs). The MB lobes can be subdivided into 15 consecutive sectors based on their innervation by distinct types of DANs and MB output neurons (MBONs). Evidence suggests that the unconditioned aversive stimulus is relayed via three types of dopaminergic neurons innervating the α’1γ2, β2β’2, and γ1 sectors, respectively, thereby modulating synaptic strength between KCs and MBONs and leading to conditioned behaviours^4,5,26–29^. Multiple sessions of training with regular rest intervals (i.e., spaced training) induce a transient increase in KC-MBON responses that eventually transforms into stable LTM—a process that involves local protein synthesis in three types of MBONs innervating the α3/β′1/β′2/γ3 sectors, respectively^4,5^. It remains unclear how this sector-specific modulation occurs and leads to LTM storage in specific neurons and synapses downstream of the MB.

The dorsal paired medial (DPM) neuron, on the other hand, is a single giant neuron with extensive neurites that innervate all MB lobes, modulating KC-MBON activity by releasing serotonin to sustain neural activity associated with an intermediate phase of anesthesia-resistant memory (ARM)^27^. In aged flies, reduced DPM-MBON connectivity appears to impair protein synthesis-dependent LTM^28^. Thus, ARM and LTM may be mutually exclusive^28^. To address how DPM participates in memory formation, we first trained flies with DPM serotonin levels reduced by adult-stage specific RNAi-mediated downregulation of synthetic enzymes. Normally, flies form maximal 1-day memory after 10 sessions of spaced training (10x spaced) but minimal memory after only three sessions of spaced training (3x spaced). When serotonin signaling in the DPM neuron was reduced, 3x spaced training now was sufficient to produce maximal 1-day memory (**Fig. 3a**). This enhanced LTM lasted for at least 4 days and was not seen after 3x massed training (**Fig. 3a**), suggesting a bona-fide protein-synthesis dependent LTM (and not ARM). With immunolabeling, we also observed a significant increase in vesicular monoamine transporter (VMAT) proteins—which uptake serotonin and other monoamines into presynaptic vesicles—at the DPM soma within 3 hours after 3x spaced training (**Fig. 3b**). This increase in VMAT expression was abolished by acute activation of temperature-sensitive RICIN^cs^, a ribosomal toxin that inhibits protein synthesis^3,30,31^. RNAi-mediated downregulation of VMAT yielded a decrease in anti-VMAT intensity in the DPM soma (**Fig. 3c**) and also enhanced 1-day memory after 3x spaced training (**Fig. 3d**). Together, these results suggest that insufficient spaced training suppresses LTM formation by inducing VMAT synthesis and increasing serotonergic signaling from DPM neurons.

**Fig. 3.**
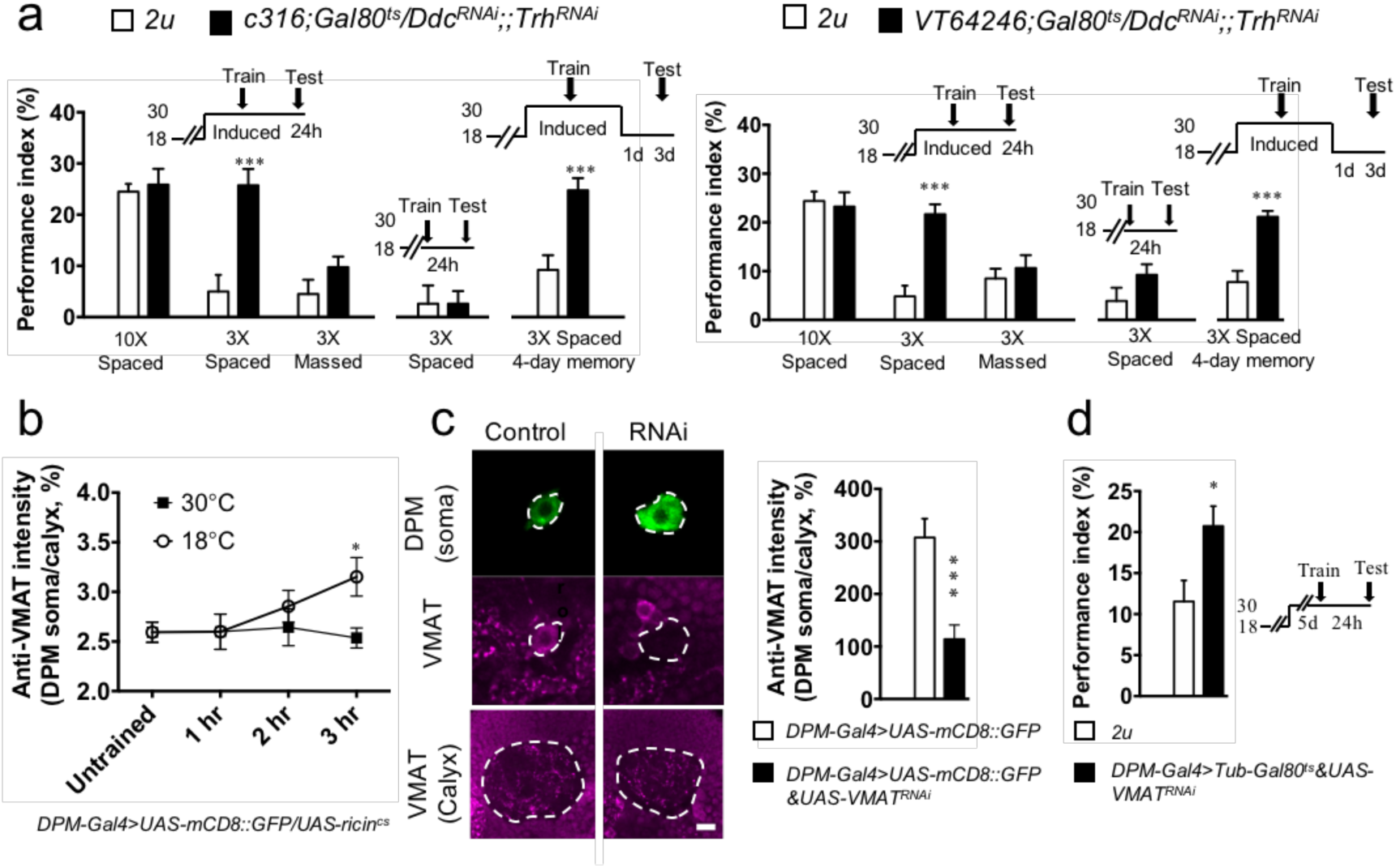
Serotonin released from dorsal paired medial (DPM) neurons suppresses long-term memory (LTM) formation. **(a)** Effects of adult-stage specific down-regulation of serotonin synthesis enzymes (*Ddc^RNAi^* and *Trh^RNAi^*) with two independent *DPM-Gal4* drivers (*c316* and *VT64246*) on 1-day and 4-day memory retention after various training protocols. **(b)** Changes in VMAT signals in the DPM soma within 3 hours after training. Protein synthesis was blocked by the active ribosomal toxin RICIN^cs^ at 30°C but unaffected by the inactive RICIN^cs^ at 18°C. **(c)** Effectiveness and specificity. Left, *VMAT^RNAi^* (vesicular monoamine transporter) effectively downregulates anti-VMAT immuno-positive signals (magenta) in the DPM cell body (green) but not in the mushroom body (MB) calyx, which is not innervated by the DPM neuron. Right, Quantitative measurements. **(d)** Adult-stage specific RNAi-mediated downregulation of VMAT in the DPM neuron enhanced 1-day memory (*DPM-Gal4*: *VT64246-Gal4*) (see **Methods** for training protocols). Each value represents the mean ± S.E.M. (n = 8 in **A, B, D,** n = 10 in **C**). *, P < 0.05; ***, P < 0.001.

### Visualizing synaptic VMAT molecules in DPM neurites

To further examine how insufficient training-induced VMAT proteins regulate serotonin release from DPM neurons in synapses, we used single-molecule DTLM imaging to quantify changes in VMAT distribution among all DPM neurites before and after 3x spaced training (**Fig. 4a**). By cropping the MB boundary, we determined the total number of immunolabeled VMAT molecules in the MB. We classified these VMAT molecules into DPM+ and DPM-groups based on the 3D digital intersection between VMAT and DPM (**Supplementary Movie 4**). Importantly, this classification can be achieved only by using DTLM **(Supplementary Fig. 7a)** and cannot be achieved with state-of-the-art confocal microscopy **(Supplementary Fig. 7b)**. The precision of DPM+ VMAT localization was demonstrated by targeted *VMAT^RNAi^* expression, which reduced the total number of VMAT molecules in DPM+ neurites but not in the DPM- regions of MB lobes **(Fig. 4b, Supplementary Fig. 7c**). Next, we examined VMAT molecules throughout the DPM neurites before and 3 hours after 3x spaced training and found that their distributions are highly variable among different flies, with a tendency to increase after training **(Supplementary Fig. 7d).** Unexpectedly, this tendency was statistically insignificant (**Fig. 4b**), regardless of the increase in the soma (**Fig. 3b**). Connectomic electron microscopy (EM) tracing of the α1/α2/α3 MB sectors indicates that the DPM neuron synapses with DANs/KCs/MBONs to form intricate sector-specific networks^32,33^. These DPM neurites exhibit branch-specific neural activity (memory traces) after associative learning^34^, prompting us to investigate whether 3x spaced training induced VMAT expression in some but not all sectors of DPM neurites.

**Fig. 4.**
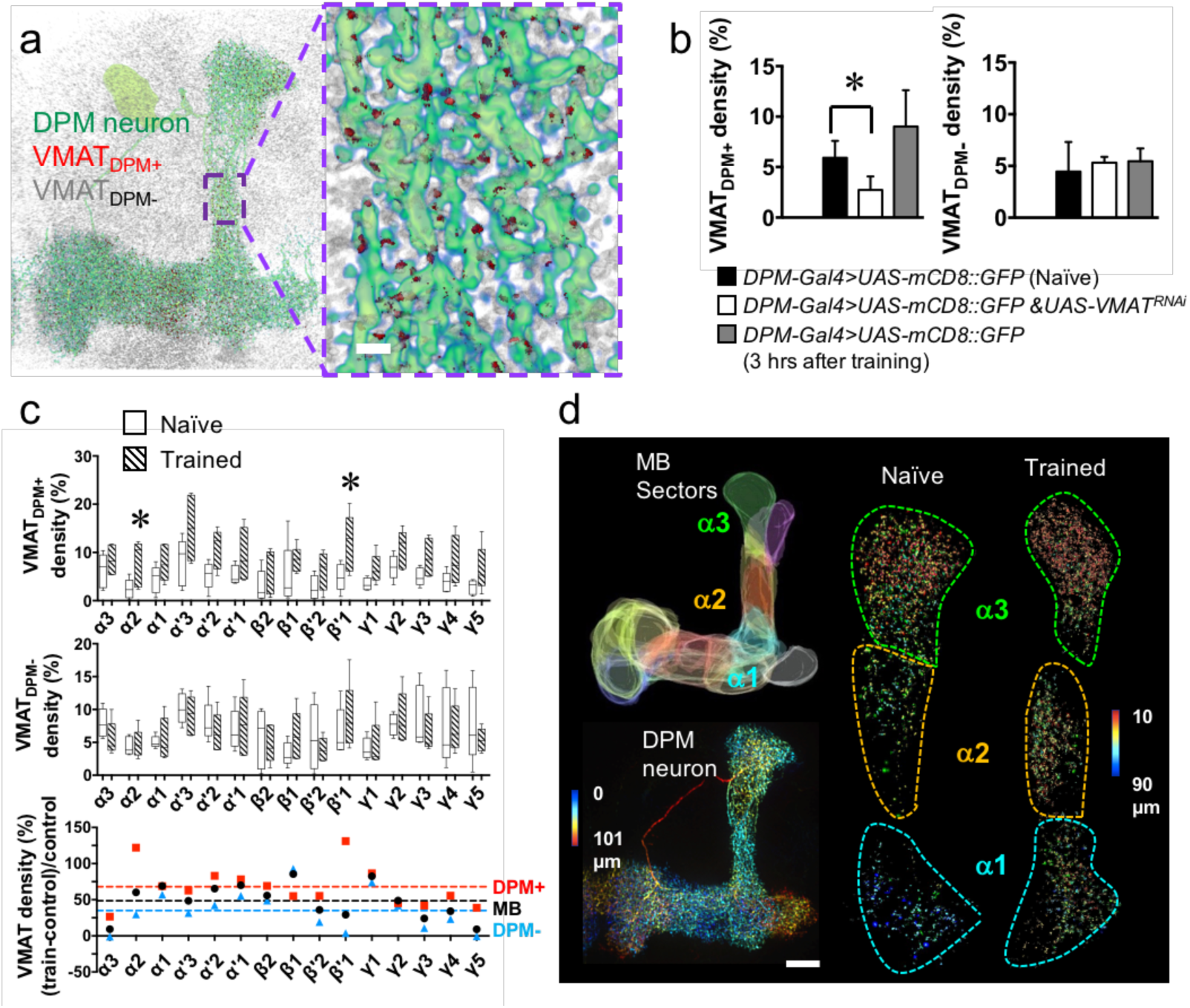
Training induces sector-specific increases in VMAT expression in axons of DPM neurons. **(a)** Visualization of VMAT molecules. Left, VMAT proteins (gray) distributed within (red) or outside of (gray) the DPM neurons (green). Right, an enlarged view. **(b)** *VMAT^RNAi^*-mediated changes in VMAT density in the DPM+ neurites (left) and DPM- regions (right) within the MB 3 hours after 3x spaced training. **(c)** Comparison of VMAT distribution in DPM axons between naïve and trained flies. VMAT density increases >100% in the α2 and β′1 sectors of DPM+ neurites (red) but not DPM- regions (blue) after 3x spaced training. The dashed line indicates the average change among different sectors [Each value represents the mean ± S.E.M. (n = 6 in **B**; n = 5 in **C**); * = P < 0.05; see **Supplementary Fig**. 8 for calculations]. **(d)** Upper left, a schematic representation of MB sectors. Lower left, a single DPM neuron innervates all MB lobes. Right, a representative volume image shows the training-induced increase in the number of VMAT proteins in the DPM neurites innervating the α2 sector. Color code indicates depth.

### MB sector-specific increases in DPM VMAT molecules induced by space training

We manually segmented DPM neurites into 15 MB sectors based on methods utilized in a previous study^35^ (**Supplementary Fig. 8 and Methods**). Quantitative analysis revealed that the density of VMAT molecules in DPM+ neurites increased after training in the α2 and β’1 sectors but not in the remaining MB sectors or in DPM- regions, when compared to that in naïve flies **(Fig. 4c, d**). These increases appeared to be evenly distributed in the affected sectors. In naive flies that had not undergone training, the distribution of VMAT molecules was highly variable between DPM+ and DPM- neurites and among different sectors (**Fig. 4c**).

## Discussion

Spatiotemporal allocation of associative learning-induced VMAT molecules suggests that DPM neurons regulate memory formation via serotonergic modulation of DAN/KC/MBON circuits in specific MB sectors. Translational regulation may account for this sector-specific allocation, as increases in VMAT are (i) regulated by Fragile X mental retardation protein, which itself is involved in olfactory LTM formation^36,37^ and (ii) blocked by inhibition of protein synthesis (**Fig. 3d**). Transcriptional regulation may also be implicated, as increases in VMAT expression after training occur in both the soma and neurites of DPM neurons (**Figs. 3** **and 4**). Previous studies have suggested that training induces local protein synthesis in α3/β′1/β′2/γ3 MBONs during LTM consolidation^4,5^. The findings of the present study suggest that modulation of serotonergic signaling (via VMAT) from DPM neurons to sector-specific MB neurons also contributes to LTM consolidation during the first few hours after training (**Fig. 3d**) but not during LTM retrieval^38^. Further studies are required to determine whether α2/β′1-VMATs in the DPM directly and specifically modulate the function of α2/β′1 MBONs. Although α2 MBONs exhibit highly variable functional responses among individual subjects^39^, their outputs appear necessary during LTM retrieval^39^. In contrast, β′1 MBONs appear to be involved in the forgetting process^40^.

Neural plasticity involves local protein synthesis in both presynaptic axon terminals and postsynaptic dendrite spines^41^. In *Drosophila* olfactory memory formation, postsynaptic regulation of protein synthesis has been demonstrated^4,5^, and here we present data suggesting presynaptic regulation of protein synthesis during memory formation, as well. Using deep-tissue localization microscopy, we further extend the visualization of protein allocations in a few target neurons to a systems level of analysis. Consistent with branch-specific functional responses during associative learning^34^, our finding of sector-specific allocation of presynaptic VMAT proteins in DPM neurons adds another layer of coding complexity to the DANs/KCs/MBONs circuit within the MB. Spatiotemporal VMAT-mediated serotonin release from DPM axons then directly modulates the neural activity of a subset of KCs within specific MB sectors. Such serotonergic signaling helps to translate transient neural activity in the MB circuit during early memory into LTM that includes postsynaptic regulation of protein synthesis in specific MBONs.

One limitation of our current imaging method is that the precision of axial localization is restricted by the thickness of the Bessel lightsheet (~500 nm), which is much less than the precision at the lateral plane (~30 nm). Thus, separating entangled neurites is trivial at X-Y plane but not always reliable along the Z axis. In this study, we corrected for this Z-plane limitation by classifying VMAT immune-positive signals into DPM+ versus DPM- groups. This approach was reasonably reliable because all imaged neurites in the MB derived from a single DPM neuron. In future studies, the Z-axis resolution may be improved further (i) by optical clearing using a medium with a higher refractive index^42^ matched with a higher-aperture objective, (ii) by using a cylindrical lens to refine the Z position^43^, and/or (iii) by combining DTLM with expansion microscopy^44^. With imaging near isotropic super-resolution, visualization of the connectome representing LTM-dependent neuroplasticity among all synaptic connections in an intact fly brain soon will be achievable^3–5^.

## Acknowledgments

We are grateful to Tim Tully for critical comments and editing on the manuscript. The control software of the microscope is licensed by Howard Hughes Medical Institute, Janelia Farm Research Campus. We thank David Krantz for sharing the rabbit-anti VMAT antibody. We thank Vienna *Drosophila* Resource Center, *Drosophila* Genetic Resource Consortium and Bloomington Stock Center for fly stocks. This work was supported by grants from the Ministry of Science and Technology and Ministry of Education of Taiwan for L.A.C (MOST 106-2321-B-007-008-MY3), A.S.C. (Higher-Education Deep Cultivation Program) and B.C.C. (MOST 107-3017-F-007-004, MOST 103-2113-M-001-003-MY2), and from the Academia Sinica for B.C.C. (Career Development Award).

## Author contributions

L.A.C. and C.H.L. planned and performed the imaging experiments and image processing. L.A.C. performed immunostaining experiments. C.H.L, Y.T.L, and B.C.C. designed and constructed the DTLM. S.M.Y. produced the HMSiR conjugates. K.L.F. and C.C.C. planned and performed the behavioral experiments. Y.T.L. designed the automated blinking image processing protocol and software. L.A.C. performed image analysis. L.A.C., C.H.L., B.C.C. and A.S.C. wrote the manuscript. B.C.C and A.S.C. planned and managed the project.

## Data and materials availability

Requests for materials should be addressed to B.C.C. or A.S.C.

## Methods

### Fly stocks

Fly stocks were raised on cornmeal food at a temperature of 25°C and relative humidity of 70% under a 12-h light/dark cycle. The following fly lines were used in the current study: *Fruitless-Gal4* (66696, Bloomington *Drosophila* Stock Center) was used to label neck neurons*, MZ19-Gal4* (34497, Bloomington *Drosophila* Stock Center) was used to label olfactory projection neurons, *12862-Gal4* (111501, DGRC) was used to label giant fiber neurons, *TH-Gal4* (8848, Bloomington *Drosophila* Stock Center) was used to label dopaminergic neurons, *c316-Gal4* (30830, Bloomington *Drosophila* Stock Center) and *VT64246-Gal4* (v204311, VDRC) were used to label DPM neurons, *UAS-mCD8::GFP* (5137 and 5310, Bloomington *Drosophila* Stock Center)and *UAS-GCaMP6f* (42747, Bloomington *Drosophila* Stock Center) were used as reporters of Gal4 expression, *UAS-Dscam[1.7]::GFP* (From T. Lee, Howard Hughes Medical Institute, Ashburn, VA) was used to label Dscam in Gal4-labelled neurons, *tub-Gal80^ts^* (From L. Luo, Stanford University, Stanford, CA) was used to block Gal4 expression at 18°C, *UAS-Ddc^RNAi^* (3329, VDRC) and *UAS-Trh^RNAi^* (35240, VDRC) were used to downregulate serotonin expression in DPM neurons, and *UAS-VMAT RNAi* (v104072 and v4856, VDRC) was used to downregulate VMAT protein expression in DPM neurons.

### Immunohistochemistry

Fly brains were dissected in PBS (pH 7.2) and immediately transferred to a microwave-safe 24-well plate containing 4% paraformaldehyde in PBS. The plate was placed on a shaker for 25 min. Fixed tissues were then permeabilized and blocked in PBS containing 2% Triton X-100 and 10% normal goat serum (NGS; Vector Laboratories, Burlingame, CA) at 4°C overnight. Immunostaining was sequentially performed in PBS containing 1% Triton X-100 and 0.25% NGS using the following primary antibodies: mouse anti-Discs large antibodies (antibody 4F3; 1:20 dilution; Developmental Studies Hybridoma Bank, Iowa City, IA), rabbit anti-GFP antibodies (1:250 dilution; Thermo Fisher Scientific Inc, A11122), rabbit anti-VMAT (1:250 dilution), and secondary antibodies including either biotinylated goat anti-rabbit immunoglobulin G (IgG) (1:250 dilutions; Thermo Fisher Scientific Inc). Biotin-conjugated IgG was detected using Alexa Fluor streptavidin 635 (1:500 dilution; Thermo Fisher Scientific Inc) or HMSiR streptavidin (1:1000 dilution). Each step was carried out over the course of 2 days, with extensive washes between steps at room temperature (25°C). Samples were then transferred to ScaleView-A2 for 2 days before imaging.

### HMSiR conjugates

For goat anti-rabbit labelling of HMSiR, we incubated 250 µg of goat anti-rabbit F(ab’)2 fragment (Jackson Immunoresearch, West Grove, PA) in 0.1 M sodium borate buffer at pH 8.5 with 0.8 µl of 10 mM HMSiR-NHS (GORYO Chemical, Bunkyo-ku, Tokyo) in DMSO at 37°C for 30 min, after which the sample was incubated at 4°C overnight. The conjugated antibody was then passed through a Desalt Z-25 column (emp Biotech GmbH, Berlin) to remove unconjugated HMSiR-NHS, and the medium was replaced with PBS (pH 7.2). To label HMSiR with streptavidin, we incubated 125 µg of streptavidin (Sigma-Aldrich, St. Louis, MO) in 250 µl PBS at pH 7.2 with 0.8 µl of 10 mM HMSiR-NHS in DMSO, after which we utilized the same protocol as that used for antibody labelling. Concentrations were then measured at an absorbance of 280 nm, after which the HMSiR conjugates were stored at 4°C until use.

### Microscope optics and image acquisition

The lasers were combined with long-pass dichroic filters and aligned collinearly before entering an acousto-optical tunable filter (AOTF), which is used to control laser exposure time and wavelength. The laser beam passing through the AOTF was expanded to 4 mm full width at half maximum (FWHM) to distribute the energy evenly onto the annular ring pattern at the quartz mask for creating the illumination pattern conjugated to the back focal plane of the objective lens. To conjugate the illumination pattern to the scanning mirrors, a pair of lenses aligned in 4F geometry was inserted between the mask and scanning mirrors (L12, L13, **Fig. 1a**). Another 4F lens pair was inserted between the galvo mirrors to relay the pattern (L14, L15**, Fig. 1a**). After reaching the scanning mirror (Cambridge Technology, 6215H), the pattern was magnified and conjugated to the back aperture of the excitation objective through another pair of lenses (L16, L17, **Fig. 1a**). A dip-in objective was placed perpendicular to the illumination plane to collect the fluorescence signal. Using a piezo scanner (Physik Instrumente, P-725.4 PIFOC), the detection objective was moved in synchrony with the position of the lightsheet, which was controlled by the scanning mirror. Between the individual volumetric scans, an additional settle time on the order of milliseconds was included to stabilize the piezo scanner.

The length of the self-reconstructing axial extent of the Gaussian-Bessel lightsheet can be controlled by the geometry of the illumination pattern^16^. A ring-shaped illumination will transform into a concentric irradiance profile distributed along the radial direction around the optical axis at the focal plane of the excitation objective lens. The illumination profile and the total power carried by the Bessel beam are controlled by the diameter and thickness of the annular ring. In the present study, the geometry of the illumination pattern was chosen based on a balance between the area of coverage and the power density of the lightsheet. To achieve stochastic blinking with reasonable signal to noise ratio (SNR), it is necessary to fill the observation plane with a power density above 40W/cm^2^. To localize subareas within the fruit fly brain, we used a lightsheet with an axial FWHM of 50 μm (outer N.A. = 0.26, inner N.A. = 0.185).

As the size of the observation area increases, however, the power density of the lightsheet created by simply filtering the laser profile using an annular ring mask is insufficient for providing high precision localization because of the decreasing SNR, as the annular ring intrinsically filters out the most intense component from the Gaussian irradiance profile (**Fig. 1d2, e1**). In this optical configuration, the field-of-view (FOV) cannot be extended further. We therefore used an axicon lens (Thorlabs) to concentrate the laser energy into a ring pattern, following which the unwanted components were filtered using an annular ring mask (outer N.A. = 0.187, inner N.A. =0.174). The length of the lightsheet generated by the axicon lens can extend to over 200 μm to cover the entire cross section of the fruit fly brain while maintaining a sufficient power density to excite HMSiR (**Supplementary Fig. S1d3, e2**).

Choice of the objective lens pair depends on the type of immersion medium. In PBS, a customized excitation objective (Special Optics, 0.65 NA, 3.74 mm WD) was used for image acquisition, while a water immersion detection objective lens (Nikon, CFI Apo LWD 25XW, 1.1 NA, 2 mm WD) was used for signal collection. For image acquisition in ScaleView-A2, the excitation objective lens was replaced by a customized objective lens (N.A. = 0.5, working distance = 12.8 mm; NARLabs, ITRC, Taiwan, **Fig. 1b**) designed to optimize performance in media with a higher refractive index (n = 1.38). An immersion detection objective (Olympus, XLPLN25XSVMP2, 25X, 1.0 NA) was used for single-molecule detection in the high-refractive index medium. In the present study, the lightsheet system was carefully calibrated to facilitate single-molecule detection with respect to each of the experimental conditions. As shown in Extended Data Fig. 1, the point spread function (PSF) values from different system configurations were compared to quantify the image quality of the lightsheet microscope system.

Single molecule fluorescence was detected using a sCMOS camera (Hamamatsu, Orca Flash 4.0 v2 sCOMS) equipped with a tube lens (f = 500 mm). The exposure time of each frame is typically 100 ms, while the period of an individual volume stack is approximately 40 s (400 layers in one stack).

Samples were loaded onto 5-mm round glass coverslips (Warner Instruments) using 100 nl of Cell-Tak™ (Corning®). During image acquisition, samples and both objectives were immersed in a chamber to maintain optical clarity. All experiments were conducted at room temperature.

The use of a Bessel beam lightsheet allows for single-molecule localization even when tissues are maintained in PBS, although the uncertainty of molecule localization increases along with the depth of the illumination plane (**Supplementary Fig. 3a, Supplementary Movie 1**). This increases the full-width at half-maximum intensity and reduces image quality in the brain, compared to images acquired from the relatively thinner neck (**Supplementary Fig. 3b**). To achieve single-molecule localization in deep tissues, we integrated the lightsheet with optical tissue clearing technology^45,46^. When the fly brain is cleared with neutral ScaleView-A2 solution^18^, the Bessel lightsheet provides enough energy to excite a dense population of HMSiR fluorophores at high efficiency throughout the entire brain. As expected, blinking signals are much clearer in ScaleView-A2 than in PBS, especially beyond a depth of 30 μm (**Supplementary Fig. 4**). Moreover, the localization uncertainties remain invariant as the imaging depth increases (**Fig. 1d**). This single-molecule DLM imaging method allows for the three-dimensional reconstruction of olfactory projection neurons, whose axons extend from the antennal lobe at the frontal surface of the brain to the calyx at the posterior surface (**Supplementary Fig. 5a).** With over 50 million molecules localised at a lateral precision of approximately 30 nm, the quality of these DLM images represents a substantial improvement over conventional lightsheet images acquired using the same optical parameters **(Supplementary Movie 2**).

### Image processing and rendering

Localization images were reconstructed using ThunderSTORM^19^, an ImageJ plugin with a self-built macro for batch processing of massive data. As schematically depicted in **Supplementary Fig. 2**, data were initially transposed to time lapse series on a local workstation, after which they were transferred to remote Lustre storage and distributed to a three-node Torque cluster (Intel Xeon X5660 with 48 GB memory each, connected to Lustre storage). After a particle list was generated for each layer, the list was deposited into one of the clusters, after which a single list containing all localization events within the imaging volume was generated. The list was then used to render the reconstruction image for presentation and analysis. ThunderSTORM was used to perform drift correction using fiducial marker tracking or the cross-correlation method. For three-dimensional volume rendering, the image stack was resampled to minimize discontinuities in structural integrity.

We used the Material Statistics module in Avizo 9.4 (Thermo Fisher Scientific Inc) to quantify the volume of MB sectors, DPM neurons and VMAT protein expression. This module calculates the voxel numbers inside a labeled area, which can be transformed into volumes by multiplying by a known voxel size in each image. We manually segmented the boundaries of MB sectors using the Lasso tool in segmentation mode. We used the Magic Wand tool to select one seed within the DPM neuron and to determine a reasonable threshold for selecting connecting voxels. We directly used the threshold tool for whole-volume VMAT images, setting the lower bound to 78 (8-bit; 0-255).

### Analysis of image acquisition speed and resolution

Resolution in localization microscopy is governed by localization precision and the density of localization events. When resolving novel structures, an extremely high localization density is required^13^. Very long image acquisition times are required for this outcome, however, which slows experimental throughput. To determine a realistic acquisition time with a reliable statistical basis, we plotted resolution with respect to time. For localization of VMAT expression in DPM neurons, localization density (number of localization events/area of the structure) increases exponentially with acquisition time. As the number of sampling frames increases, the growth of localization density slows due to photo-bleaching and depletion of dye molecules (**Supplementary Fig. 9a)**. The theoretical resolution limit is estimated by the localization precision and the Nyquist resolution^47,48^. Resolution power increases rapidly in the first 300 frames (**Supplementary Fig. 9b)**. Afterwards, resolution power slows and finally converges at 86 nm. This represents a lower bound of structural resolution that can be achieved with our current methods.

More generally, if an infinite photon budget existed and no photo-bleaching occurred, localization density would increase monotonically with the number of sampling frames. Such modeling indicates that resolution reaches 25 nm when 10,000 frames per layer are used in reconstruction—which is 20 times the acquisition time used in this study. Our observed dependence of resolution on sampling frames reveals an exponential relation (likely due to photo-bleaching and limited photon budget; **Supplementary Fig. S9c)**, which suggests that the acquisition time required for maximal resolution may be much longer.

Fourier ring correlation (FRC) provides a quantitative determination of the structural characteristic in the reconstructed image^49^. The FRC value at the 1/7 cutoff frequency in VMAT protein localization is 283.6 ± 69.9 nm (**Supplementary Fig. 9d)**. It should be noted that FRC analysis is based on weighting of the spatial frequency domain and is dominated by the primary feature within the image. As shown in the inset of (**Supplementary Fig. 9d**), VMAT protein presents an island feature with a dimension from hundreds of nanometers to micrometer, which leads to a higher estimated FRC value.

When considering the statistics of localizing VMAT molecules, the distribution of VMAT is confined to DPM axons. Consequently, the reliability of the results is dominated by localization precision rather than structural determinations. Regardless, we kept the number of frames of time-lapse data used in VMAT localization to 400 to 500 per layer to ensure all analyses were performed based on similar localization density and theoretical resolution. Acquisition time for one VMAT dataset was approximately 5.6 hours (100 ms per frame, 401 frames per imaging volume, 500 volumes recorded). For localization of anti-TH signals (**Fig. 2**), acquisition time was approximately 5.8 hours (80 ms per frame, 521 frames per imaging volume, 500 volumes recorded). With four sub-volumes per whole brain, total acquisition time took approximately 23.2 hours to complete.

### *Drosophila* memory assay

Flies were subjected to aversive olfactory conditioning 2 to 5 days after eclosion. Prior to conditioning, flies were accommodated to a behavioral room with a temperature of 20°C and relative humidity of 70% for 30 min. At the start of olfactory conditioning, approximately 80 flies were transferred to the training tube, after which two aversive odors (3-octanol (OCT); dilution: 1.5 x 10^-3^; Sigma-Aldrich) and 4-methylcyclohexanol (MCH); dilution: 1.0 x 10^-3^; Fluka) were delivered successively in a current of air (750 ml/min) for 60 s at intervals of 45 s. The first odor (CS+) was paired with 12 pulses of electric foot shock at 65 V (serving as the unconditioned stimulus [US]), while the second odor (CS-) was not. This process represented a single session of training. In our experiment, we conducted three sessions of spaced training, with a 10-min interval between each cycle.

*Gal4* expression was inhibited by *Gal80^ts^* by maintaining flies at 18°C. One day before training, flies were moved from 18°C to 30°C, thereby deactivating *Gal80^ts^*. Control flies were maintained at a constant temperature of 18°C. Wild-type and experimental flies carrying the same transgenes were trained using several different protocols: three spaced sessions (3x spaced), three massed sessions (3x massed), or 10 spaced sessions (10x spaced).

### Statistical analysis

For behavioral experiments, control and treatment groups were tested together (balanced and blinded), with sample sizes listed in figure legends. Because performance indices were normally distributed, the significance of each treatment-versus-control paired comparison was tested using a two-tailed Student’s t-test, with P values indicated in figures.

